# Localized Chemogenetic Silencing of Inhibitory Neurons: A novel Mouse Model of Focal Cortical Seizures

**DOI:** 10.1101/2020.11.04.367862

**Authors:** Adi Miriam Goldenberg, Sarah Schmidt, Rea Mitelman, Dana Rubi Levy, Yonatan Katz, Ofer Yizhar, Heinz Beck, Ilan Lampl

**Affiliations:** Department of Neurobiology, Weizmann Institute of Science, Rehovot 7610001, Israel; Institute for Experimental Epileptology and Cognition Research, University of Bonn, 53105 Bonn, Germany

## Abstract

Focal cortical epilepsies are frequently refractory to available anticonvulsant drug therapies. One key factor contributing to this state is the limited availability of animal models that allow to reliably study focal cortical seizures and how they recruit surrounding brain areas *in-vivo*. In this study, we selectively expressed the inhibitory chemogenetic receptor, hM4D, in GABAergic neurons in focal cortical areas using viral gene transfer. Following focal silencing of GABAergic neurons by administration of Clozapine-N-Oxide (CNO), we demonstrated reliable induction of local epileptiform events in the electroencephalogram (EEG) signal of awake freely moving mice. Experiments in anesthetized mice showed consistent induction of focal seizures in two different brain regions – the barrel cortex (BC) and at the medial prefrontal cortex (mPFC). Seizures were accompanied by high frequency oscillations, a known characteristic of human focal seizures. Seizures propagated, but an analysis of seizure propagation revealed favored propagation pathways. CNO-induced epileptiform events propagated from the BC on one hemisphere to its counterpart and from the BC to the mPFC, but not vice-versa. Lastly, post-CNO epileptiform events in the BC could be triggered by sensory whisker-pad stimulation, indicating that this model, applied to sensory cortices, may be useful to study sensory-evoked seizures. Taken together, our results show that targeted chemogenetic inhibition of GABAergic neurons using hM4D can serve as a novel, versatile and reliable model of focal cortical epilepsy suitable to systematically study cortical ictogenesis in different cortical areas.

**Significance Statement:** Focal cortical epilepsies are often hard to alleviate using current anticonvulsant therapies while further drug discovery is impeded by the limited variety of suitable animal models. In this study, we established a novel model of focal cortical seizures induced by spatially-restricted chemogenetic silencing of cortical inhibitory neurons. We have shown this method to be effective at various cortical regions and reliably induce seizures that share key characteristics with known human epilepsy traits, including sensory triggering and seizure propagation. This model may thus be used to advance the discovery of new remedies for focal cortical epilepsies, as well as to improve our understanding of seizure spread along different cortical pathways.

## Introduction

Epilepsies are one of the most common neurological disorders, with ~50 million patients world-wide. Amongst the different types of epilepsies, many disease entities arising from cortical foci have proven particularly difficult to treat. A substantial fraction of >30% of these patients are refractory to currently available anticonvulsant drugs. Importantly, the success rate of epilepsy surgery is not nearly as high as that reported for non-cortical epilepsies such as temporal lobe epilepsy. Thus, cortical epilepsies remain a significant clinical problem.

In many cases, animal models of acute and chronic seizures have driven drug discovery. A large number of commonly used anticonvulsant drugs have been discovered using acute seizure models, such as the maximal electroshock or the subcutaneous pentylenetetrazol seizure model (Löscher, 2011). Subsequently, other models have been added in the framework of large drug screening programs, such as the 6Hz psychomotor seizure model. These acute induced seizure models allow excellent control over the seizure onset and are well suited to high-throughput studies of drug effects. However, they do not allow studying the effects of regional/focal ictogenesis, and do not display spontaneous seizures relying on epilepsy-related changes in the neuronal network.

The addition of chronic epilepsy models, such as pilocarpine, kainic acid and some types of kindling model allows to overcome this issue. These models present spontaneous seizures and share many similarities with human temporal lobe epilepsy (Kandratavicius et al., 2014). However, the incidence of spontaneous seizures is low, making large-scale screening of drugs extremely labor-intensive. This problem is accentuated in cortical epilepsies. Chronic models of cortical epilepsies have a low number of seizures, which are frequently difficult-to-detect. Perhaps for this reason, models of focal cortical epilepsies have been much less commonly used for drug discovery.

Acute models of focal cortical seizures include focal application of chemoconvulsants like 4-aminopyridine (Rothman, 2009; Wenzel et al., 2017). These approaches provide a way to assess cortically induced seizures, and their spread, but using application of chemicals that spread is inherently variable and does not allow to pinpoint the region and cell types of onset precisely.

Recent optogenetic and chemogenetic developments have allowed these issues to be addressed. Most of the optogenetic and chemogenetic approaches in the field of epilepsy have focused on suppressing seizures in classical epilepsy or seizure models (for a review please refer to Forcelli, 2017 and to Walker and Kullmann, 2020). Few studies have also used these methods to trigger or enhance seizures in the classical models (Krook-Magnuson et al., 2015; Sorokin et al., 2017; Chen et al., 2020). Models using opto- or chemogenetic actuators to induce epileptic seizures de-novo are scarce, and to the best of our knowledge, none has induced seizures using chemogenetic manipulation of inhibitory neurons.

In this paper, we present a novel cortical seizure model, in which epileptiform activity can be reliably induced in a restricted cortical area by chemogenetic silencing of local GABAergic neurons. This model was developed in light of the widely accepted assumption that epileptic seizures are accompanied and even caused by an increased excitation-inhibition ratio. This idea was introduced in 1997 by Schwartzkroin (Schwartzkroin, 1997), and has since been supported by ample theoretical and empirical evidence both *in vivo* and *in vitro* (Lopes da Silva et al., 1994; Bradford, 1995; McCormick and Contreras, 2001; Stief et al., 2007).

In our study, we targeted hM4D expression to either the medial prefrontal cortex (mPFC) or the barrel cortex (BC) of Gad2-IRES-Cre mice. Following the expression of the receptor in inhibitory interneurons, we recorded the network response to its synthetic ligand, Clozapine N-oxide (CNO), in awake and anesthetized mice. These experiments provided tests of CNO response propagation and quantification in high frequency bands (80-500Hz). We also examined the CNO effect on the response to sensory stimulation. All these experiments indicate that this model mimics well some of the central characteristics of the disease, suggesting it can be used as a promising new epilepsy model.

## Methods

### Animals and IACUC

Procedures involving animals were reviewed and approved by the Weizmann Institute Institutional Animal Care and Use Committee. We used 6-11 weeks old heterozygous Gad2-IRES-Cre mice (Jax #010802, The Jackson Laboratory, Bar Harbor, ME, USA) of either sex, housed up to five in a cage with a 12-hr/12-hr dark/light cycle.

### Virus injection

For stereotaxic viral injection mice were sedated with isoflurane and placed into a stereotactic frame, where deep anesthesia was maintained over the course of the surgery (1-2% isoflurane in O_2_, v/v). A small hole (<=0.3mm diameter) was made over the intended injection site, being the mPFC (AP +1.97mm, ML 0.35mm right, DV −2.44-(−2.8)mm all measured from bregma) or the BC (AP −1.3mm, ML 2.99-3.05mm right from bregma, DV −0.45-(−1.0)mm below dura).

Virus injection was carried out using a microinjection pump (UMP3, World Precision Instruments, Sarasota, FL, USA) and a glass pipette (Wiretrol II #5-000-2005, Drummond Scientific Company, Broomall, PA, USA), pulled to have an outer diameter of 60-90μm at the tip. Each mouse underwent a single injection of a Cre-dependent virus expressing both hM4D and ChR2 (E123T/T159C) (AAV2/1-hSyn-DIO-ChR2(TCET)-P2A-hM4D). The ChR2 was used for *in-vitro* testing. The injection was either 500nl injected at a rate of 50nl/min in the mPFC or 750nl injected at changing rates of 20-50nl/min at the BC. The BC injections took place in two steps, first about 50% of the total volume was expelled at 0.45-0.5mm below the dura, and then the rest was injected at 0.9-1.0mm below dura. Following each injection, the pipette was held in place for another ten minutes, which were split in half between the two injection depths in the case of the first injection in the BC. Control mice were similarly injected with a Cre-dependent GCaMP6s virus (AAV1/2-hSyn-flex-GCaMP6s-WPRE).

The incision was closed using surgical stitches with reinforcement of veterinary glue. Local analgesia and antibiotic creams were applied while the mice were given buprenorphine (0.1mg/kg) and carprofen (5mg/kg) intraperitoneally. Mice were allowed to recover at least 29 days before first CNO treatment and at least another 19 days before electrophysiological recording, to assure steady state level of expression.

Mice intended for awake recording were initially sedated with an i.p. injection of ketamine-xylazine mixture in addition to the isoflurane anesthesia maintained throughout the procedure. Craniotomies of about 1mm in diameter were made above their mPFC bilaterally and a similar virus injection took place in the mPFC on either side using a Nanofil syringe. Following the virus injection, a combined optrode and electrocorticography (ECoG) drive was lowered into either right or left mPFC, so that the electrodes were placed at the center of the injection site. Next, four micro-screws were attached to the skull: two posterior ones were used as ground and reference, and two frontal-laterals were used for ECoG. The screws were connected to a Mill-max connector with an isolated steel wire. The exposed skull was then coated with Metabond and the connector was attached using dental cement. Mice were given buprenorphine at the end of the procedure and were allowed to recover for at least eight weeks.

Mice intended for slice recording were treated similarly to awake mice (though not implanted with a drive). The injected virus solution in these cases was mixed with AAV2/1-CamKIIα-TagRFP-T in 1:1 ratio to allow targeting of pyramidal cells in the acute slice preparation.

### Prompting epileptic activity

In order to verify the used dosage of CNO did not cause a severe behavioral effect in mice intended for recording under anesthesia, these mice were injected with 5μg/gr of CNO i.p. (Tocris, Bristol, United Kingdom) several times before any recording had been made. CNO stock was dissolved in dimethyl sulfoxide (DMSO) and kept frozen (−20°C) for up to four months. The injected solution was prepared no more than a week prior to the injection by diluting the stock in 1% phosphate buffer solution (PBS) so that the final solution contained 6.67% DMSO and 1μg/μl of CNO.

The CNO injection was carried out three times, approximately one week apart, and was followed by visual monitoring of the behavioral response. Mice showing motor seizure behavior such as forelimb clonus and rearing as well as falling (stage four and above in the customary scale of behavioral assessment of seizure threshold; Morrison et al., 1996) were injected with 2μg/gr Diazepam in order to terminate the seizure. Such events were extremely rare.

### Electrophysiological recording

#### Slice preparation and patch-clamp recording

Female black mice (>7 days after injection) were deeply anesthetized with 0.3ml ketamine hydrochloride (10%; Pfizer) and 0.6ml xylazine hydrochloride (2%; Bayer) and decapitated. Transverse 300μm thick medial prefrontal cortex slices were prepared on a vibratome (Leica VT 1000S) in ice-cold preparation solution containing (in mM) 60 NaCl, 100 sucrose, 2.5 KCl, 1.25 NaH_2_PO_4_, 26 NaHCO_3_, 1 CaCl_2_, 5 MgCl_2_, and 20 D-glucose (equilibrated with 95% O_2_ and 5% CO_2_). Slices were stored in the preparation solution for 30 min at 35°C. Slices were transferred to artificial CSF (ACSF) containing (in mM) 125 NaCl, 3 KCl, 1.25 NaH_2_PO_4_, 26 NaHCO_3_, 2.6 CaCl_2_, 1.3 MgCl_2_, and 15 D-glucose (equilibrated with 95% O_2_ and 5% CO_2_) and stored at room temperature. Slices were transferred to a submerged chamber perfused with ACSF and mounted on the stage of an upright microscope (Axioscope 2; Zeiss). Pyramidal cells were visualized with infrared oblique illumination optics and water-immersion objective (60X, 0.9 NA; Olympus). Somatic whole-cell recordings of principal cells in the mPFC were obtained with a BVC-700A amplifier (Dagan). Data was filtered with a low-pass filter of 1kHz and sampled at 50kHz with a Digidata 1440A interface controlled by pClamp software (Molecular Devices). Recording electrodes were made from thick-walled borosilicate glass capillaries (GB 150F 8P; Science Products) on a vertical puller (PP-830; Narishige). Recording pipettes for whole-cell recordings had a resistance of 3–6 MΩ and were filled with (in mM) 100 CsMs, 10 HEPES-acid, 0.5 EGTA, 2 Mg-ATP, 1 Na_2_-ATP and 5 QX314 chloride (pH 7.25, 275mOSM, adjusted with sucrose to 290mOsm). All experiments were performed at 31°C. Membrane potential was corrected offline for a liquid junction potential of 18.9mV. Wash-in of CNO was used during whole-cell experiments. CNO (final concentration, 10μM) was dissolved in DMSO and drug effects were analyzed after 10 min (125ml/h) in whole-cell recordings.

Light stimulation was used in order to depolarize the GABAergic cells at the virus injection site, and so to estimate the effect of CNO. The stimulation was carried out with a laser (473nm; Omikron) using a 200μM diameter light fiber, placed about 1cm above the slice. Light pulses were triggered by pClamp software (Molecular Devices, 20ms duration, 5 stimulations at 1Hz, 2 successive sweeps were averaged).

#### Electrophysiological recording in freely moving mice

All electrophysiological recordings in awake, freely moving mice were performed using an optrode drive consisting of an electrode bundle of 8 micro-wires (25 μm diameter, straightened tungsten wires; Wiretronic Inc;, Volcano, CA), attached to an 18-pin electrical connector, concentrically arranged around an optical fiber. Extracellular signals were recorded using the Digital Lynx integrated hardware and software system (Neuralynx). The electrical signal was filtered for ECoG and local field potential (LFP) (0.1-500 Hz) and amplified using a HS-18-CNR-LED unity-gain head-stage amplifier (Neuralynx) and sampled in 2 kHz. During the recording, mice were injected i.p. with the previously described 1μg/μl CNO solution (same dose).

#### Preparation for recording under anesthesia

Mice were briefly anesthetized with isoflurane (5% in O_2_) and then injected i.p. with 5-10μl/gr urethane solution (100mg/ml) and with 5μl/gr chlorprothixene solution (0.1mg/ml). Several minutes (15-60) later, and under additional isoflurane anesthesia, lidocaine was injected s.c. on the back of the mice”s head, their skull was exposed, and the intended recording locations were marked. A custom 3D printed plastic head bar commonly used in our lab (Katz et al., 2019) was attached to the skull using dental cement (RelyX, 3M). The brain surface at the intended recording sites was exposed using a 0.3 mm biopsy tool for BC recordings and 0.5 mm biopsy tool for mPFC recordings. In addition, a 27G needle attached to a saline syringe with a thin rubber tube was inserted and fastened to the mice”s stomachs with veterinary glue in order for it to later serve as an i.p. catheter.

#### Recording under anesthesia

The recording was done under urethane and chlorprothixene anesthesia while monitoring heart rate and body temperature, and after at least twenty minutes without isoflurane exposure.

Recordings were made using custom-made electrode arrays with 2-3 formvar coated nichrome electrodes each (0.001” bare diameter, 0.0015” coated), coupled to an Intan RHD2132 headstage. Signals were sampled at 10kHz using an Open Ephys acquisition board. Each array was designed such that while the main electrode was at the intended recording target, being 450μm (BC) or 1800-1900μm (mPFC) perpendicularly below the brain surface, an additional electrode was placed at about 200μm above the surface. This external electrode as well as a local grounding silver wire positioned above each recording site, were both kept in an artificial cerebrospinal fluid solution (containing: NaCl-124mM, NaHCO_3_-26mM, glucose-10mM, KCl-3mM, KH_2_PO_4_-24mM, MgSO_4_-1.3mM and CaCl_2_-2.4mM). This was done in light of a growing understanding of the extent to which volume conductance can affect the recording (Parabucki and Lampl, 2017). Arrays used for mPFC recordings were often coupled with a multi-mode optical fiber (NA = 0.39, 200μm core) positioned at about 500μm above the main electrode.

Each mouse had signals recorded at two sites simultaneously, (1) the virus injection site, which was either the right mPFC or the right BC (rBC), and (2) one additional remote site (non-injected) which was either the right mPFC, the rBC or the left BC (lBC). Signals were recorded at both sites before and after the administration of 5μg/gr mouse CNO injected through the i.p. catheter (same stock, further diluted to 0.5μg/μl).

#### Sensory stimulation

We assessed the response to sensory stimulation by placing a blunt needle attached to a galvanometer on the left whisker pad of some of the mice (all injected at the right hemisphere) before the beginning of the recording under anesthesia. For each mouse, the strength of stimulation was chosen so that it induced a noticeable response in the contralateral BC before CNO was introduced. The whisker pad stimulation protocol was given once every 5 seconds for at least 22 repetitions. The protocol included either (1) a single galvo stimulation, (2) three stimulations given at 1Hz, (3) three stimulations given at 3Hz, or (4) six stimulations given at 6Hz. For all the mice whose results are reported in this paper, the first protocol was presented both before the baseline recording and after 45 minutes or more since the CNO injection. However, repeated stimulation protocols may have only been presented after the CNO injection.

### Histology and visualization

Upon recording termination of all of the mice recorded under anesthesia, the electrodes were replaced with a heavily diI covered electrodes and reinserted at the same location for at least 3 minutes during which the mice were heavily sedated with isoflurane and a lethal dose of penthal was injected i.p.

Mice were then perfused transcardially with 1x PBS followed by 2.5% paraformaldehyde (PFA). The brain was postfixed in PFA at 4°C for additional 2 days or more (at least one of which before being dissected out of the skull), and was later transferred into 30% sucrose dissolved in 1x PBS (at 4°C). Once setting, the brain was sliced frozen to 30 μm thick coronal sections using a sliding microtome (Leica SM 2010R; Nussloch, Eisfeld, Germany) into eight wells in a serial manner, resulting with each well containing a series of sections spaced 240 μm from one another.

At least one such series of sections was stained per mouse, floating, in a procedure comprised of four steps. The sections were first put in a blocking solution (20% normal horse serum; CN-VE-S-2000, Vector Labs, Burlingame, Ca, USA) for two hours, in order to prevent nonspecific staining. In the second step the sections were incubated overnight in a solution containing primary antibodies against P2A, a short protein sequence expressed in the injected virus (1:400 rabbit anti P2A; ABS31, Merck, Darmstadt, Germany). In the third step, following PBS washing of the sections, they were incubated in a solution containing secondary antibody conjugated with a green fluorophore for about 25 minutes (1:100 goat anti rabbit conjugated to cy2: 711-225-152, ENCO, Petach Tikvah, Israel). Lastly, the sections were washed and incubated in a DNA staining solution (1:10,000 Hoechst 33342, Thermo Scientific, Rockford, IL, USA) for 3-6 minutes. Sections taken from control mice (expressing GCaMP6s) were only treated with the DNA staining solution. All steps were done at room temperature while gentle agitation of the sections. Sections were then washed, mounted on slides and covered.

Immunostained brain slices were imaged with a confocal scanning microscope (VS120-S6-W, Olympus) using a 2x objective for overview images (NA 0.08; Olympus) and a 20x objective (NA 0.75; Olympus) for injection site reconstruction (Fig. 1A-B).

**Figure 1.**
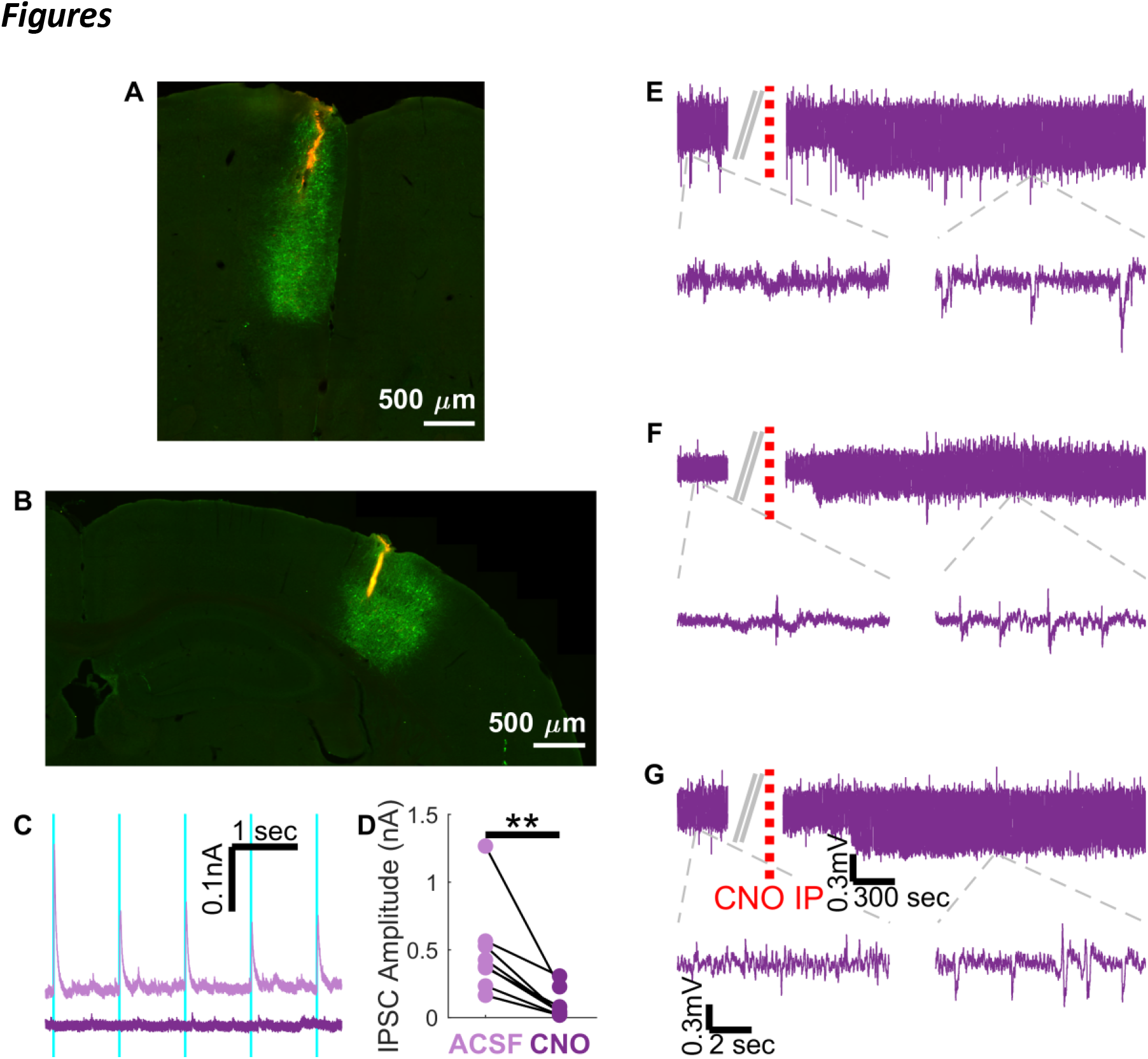
hM4D was expressed and operated as expected, and when combined with CNO, induced epileptiform events in the ECoG of awake mice. **A-B.** Immunohisto-chemistry examples demonstrating hM4D virus expression (using anti p2a staining, in green) alongside the recording electrode position (in red) in a mouse injected at the mPFC (A) or the BC (B). The hM4D expression in the mPFC injected mouse was mostly restricted to the mPFC on the medial-lateral axis though it expanded dorsally to adjacent areas. In the BC injected mouse, the expression was mostly restricted to L2-3 and L5-6, and spanned across about two or three barrels. In both cases the location of the recording electrode array coincided with the expression site. **C.** Examples of IPSCs recorded *in-vitro* in response to blue light stimulation (cyan) before (top) and during (bottom) CNO wash. The recording was done in the mPFC at an acute slice taken from a mouse expressing ChR2 and hM4D in the GABAergic mPFC neurons. In this example, CNO consistently silenced the IPSC response to ChR2 activation. **D.** Average IPSC amplitude at acute slices in response to the first ChR2 light activation before and during CNO wash across mice (N=8). CNO reduced the IPSC amplitude of each of the mice, causing a statistically significant decrease in amplitude (p=0.004). **E-G.** Top panels show examples of ECoG recordings done in three freely moving mice expressing hM4D in their mPFC bilaterally, before and after CNO administration (indicated by the dashed red lines). Bottom panels show a zoom in of the signal. The top and the bottom panels in A and B share the size bars indicated in C. All three mice show an increase in the amplitude and frequency of ECoG activity within a few minutes from the CNO injection.

### Data Analysis

All data analysis was done with custom-made software on Matlab (Mathworks). Statistical testing was done using either Wilcoxon signed-rank test (for paired measurements) or Wilcoxon rank sum test (for un-paired measurements) with Bonferroni correction for repeated measurements when such were made (α=0.05). Unless otherwise mentioned, any analysis of a parameter across mice was done by averaging per mouse followed by a comparison of the averages across mice.

#### Analysis of acute slice recordings

IPSC amplitudes and seizure events in slice recordings were analyzed with clampfit (Molecular devices). For analysis of IPSC amplitude, traces were averaged.

#### Analysis of recordings from freely moving mice

To minimize external noise and obtain a single channel of data per CNO injection, the two ECoG channels were subtracted from each other. Next the signal was lowpass filtered below 10 Hz (four poles butterworth filter).

#### Preparation of recordings from anesthetized mice

LFP was extracted from the signal by applying butterworth lowpass filter (<50Hz, order = 2) followed by downsampling to 1kHz. In addition, in order to evaluate the activity at high frequencies, the signal of the external electrode above the brain surface was subtracted from the signal at the recording site. The result was bandpass filtered between 55 and 1000Hz using butterworth filter (order = 7), notched filtered to reduce 50Hz harmonies noise, and downsampled to 5kHz.

From each mouse, a ten-minute period that ended at about one minute before the CNO injection was used as the baseline signal for analysis purposes. Following that injection, up to 45 minutes were analyzed in order to detect when the CNO effect started and reached its maximum.

In order to assess effect onset, post CNO LFP was binned into one-minute long windows with a 15-second overlap between consecutive windows. Since epileptiform events in LFP recordings manifested as rapid negative excursions, the amplitudes of local minima which were at least 200ms apart and in the lowest 5-percentile of the overall baseline signal were compared for each window against baseline minima following the same criteria. CNO effect onset was declared at the beginning of a four or more sequence of windows with significantly bigger amplitudes than baseline local minima (using Wilcoxon rank sum test).

The ten-minute long post CNO period used for analysis was chosen such that it had the maximal average standard deviation calculated in one-minute long non-overlapping windows of the LFP signal following CNO. The middle of this analysis window was also referred to as the time in which the CNO reached its maximal effect. Any further analysis was done on the ten-minute long periods chosen to represent baseline and post CNO activity.

#### Analysis over the entire ten-minute period of baseline and post CNO signals

Power spectrum density (PSD) was calculated using Welch’s power spectral density estimate over the entire analysis period and normalized per recording site per mouse such that the maximum PSD during baseline would be one. In order to assess changes related to high frequency oscillations (HFO), PSD of the 55-1000Hz filtered signal during the analysis periods was averaged in the range of 80-250Hz (referred to as ripples) and in the range of 250-500Hz (referred to as fast-ripples).

The cross correlation between the hM4D expression site and the remote site recorded was calculated over the entire analysis period such that a high cross correlation at a positive delay represents a lag of the remote signal after the signal at the hM4D expressing site.

#### Event centered analysis

Epileptic events are manifested as negative voltage excursion. Therefore, detection of the LFP epileptic discharges was done by searching for local minima which crossed the threshold of four median absolute deviations from the median voltage during the entire analysis period (of baseline and post CNO respectively). Median based detection was chosen over other variability measurements since it is less affected by extreme data points expected in epileptic recording (Bedeeuzzaman et al., 2011; Feltane et al., 2013). In order to exclude some of the falsely detected events, minimal peak separation for detection was set to be 200ms, with the peak being more negative than the average voltage recorded 300-200ms beforehand. We also omitted events detected very near the edges of the analysis window, or near a time when aCSF was added above either of the recording sites.

Detected events were aligned in one of two ways, by peak or by onset (regarded as the time in which 10% of the amplitude was reached), both after subtracting the average voltage 300-200ms before the event peak. Event amplitude was taken as the voltage at the peak after this subtraction, while the half width of the event was calculated as the time spent below half of the amplitude around the peak.

In light of the shape of the LFP at the remote recording site during an event detected following CNO at the hM4D expression site, the peak activity at the mPFC and the lBC of rBC injected mice was considered as the maximum voltage between 50ms prior to the rBC peak and 150ms after it. In the case of mPFC injected mice the BC peak was regarded at the minimum voltage at this time range. Quantification of the timing of the remote peak across mice was done by calculating a 45 bins probability histogram between the above-mentioned time limits per mouse and averaging the histograms across mice. Inspection of the remote peak amplitudes was done by z-scoring the amplitudes at the remote site and at the hM4D expression site separately and plotting them across mice. Despite that, the correlation coefficient between the amplitudes at these sites was calculated over the pooled events across mice without the z scoring, in order to prevent trend masking.

In addition, we analyzed the time vs frequency components of the detected events using wavelet analysis. We calculated the wavelets over 600ms long segments of the high bandpass filtered signal at the hM4D expression site taken at ±300ms around each detected LFP event peak. For each event, we calculated the average of the wavelet across time in the ripples frequencies and in the fast ripples frequencies, and extracted the maximum magnitude of this average as well as its timing. For each mouse the average and the standard deviation of these two parameters was calculated and compared across mice. In order to plot the wavelet’s average across mice we normalized the average wavelet per mouse such that the maximal values in the average baseline wavelet was one.

#### Granger Causality of event propagation

In order to assess the causality in the propagation of the LFP events, ±300ms around the peak of the detected event at the hM4D expression site were taken from the LFP of both recording sites and checked whether one significantly Granger caused the other with a maximum lag of 200ms using a dedicated Matlab function (Lutz, 2020). The percentage of events showing significant causality above 0.05 alpha was compared before and after CNO across mice.

#### Analysis of sensory-evoked potentials

For each mouse, whose response to a single whisker pad stimulation was recorded both before and after CNO, we averaged the response across trials. In the case of one of the control mice where no such recording was done after CNO, the response to the first of three stimuli given at 1Hz was used as such. The average response was aligned by stimulus onset and averaged again across mice. A similar analysis was done when the response to repeated stimulation following CNO was also recorded.

## Results

We selectively expressed the inhibitory DREADD (designer receptor exclusively activated by designer drug) hM4D along with ChR2 in cortical inhibitory neurons using viral gene transfer with a single bicistronic rAAV (AAV2/1-hSyn-DIO-ChR2(TCET)-P2A-hM4D, see Methods). Histological reconstructions verified hM4D expression throughout cortical depth at the recording sites of anesthetized mice injected at either the mPFC (Fig. 1A) or the BC (Fig. 1B). Recordings from mPFC slices were used to confirm that CNO reduces inhibition at the hM4D expression site (Fig. 1C-D). The co-expression of both ChR2 and hM4D allowed us to record light-evoked IPSCs from mPFC pyramidal neurons. We found that these IPSCs were strongly diminished following application of CNO (17.5%±2.0%, n=8), indicating that hM4D was expressed in inhibitory cells and that their activity was suppressed by CNO.

In order to find if silencing inhibitory cells in the restricted injected areas is sufficient to induce seizures in awake mice, we recorded ECoG in freely moving mice bilaterally expressing hM4D in the mPFC. Examples from three mice showed indeed that silencing the inhibitory cells caused an increase in the amplitude and/or frequency of high amplitude ECoG events in a seizure-resembled manner (Fig. 1E-G). In agreement with other studies utilizing systemic injection of CNO for silencing experiments, a delay of a few minutes was found in the appearance of the epileptic events after CNO injection.

To further quantify and characterize the features of the suggested paradigm in a highly controlled manner, we analyzed LFP signals recorded in seven anesthetized mice expressing hM4D at their right mPFC and eleven anesthetized mice expressing hM4d at their right BC (rBC). Electrophysiological recordings at the virus injection site were made under urethane and chlorprothixene anesthesia at least 47 days following the virus injection (Fig. 2A), and after three CNO treatments had been given to either mouse to ensure that CNO has no severe effect in awake mice. With the exception of a single mouse who was injected in the mPFC and showed extremely minimal brain activity and abnormally low heart rate throughout the experiment (and was therefore excluded from any further analysis), all of the mice showed an increase in the size and/or frequency of epileptiform events following CNO administration (Fig. 2B-C). These events were significantly larger in amplitude and shorter in their half width in comparison to events detected during baseline using similar detection criteria, and caused a significant increase of the overall PSD (Fig. 2D-F), and for the BC particularly in the delta frequency range (Fig. 2Fi). The CNO effect was analytically detectable (see Methods) within five to 25 minutes in all animals, with a tendency to take longer to influence in mPFC injected mice, and was maximal towards the end of the analyzed period (after about 40 minutes) in many of the mice (Fig. 2G-H). Importantly, we were unable to detect a significant change in LFP activity following CNO injection in control mice lacking the expression of hM4D (Fig. 3). Hence, we conclude that restricted silencing of inhibitory cells promotes local epileptic activity.

**Figure 2.**
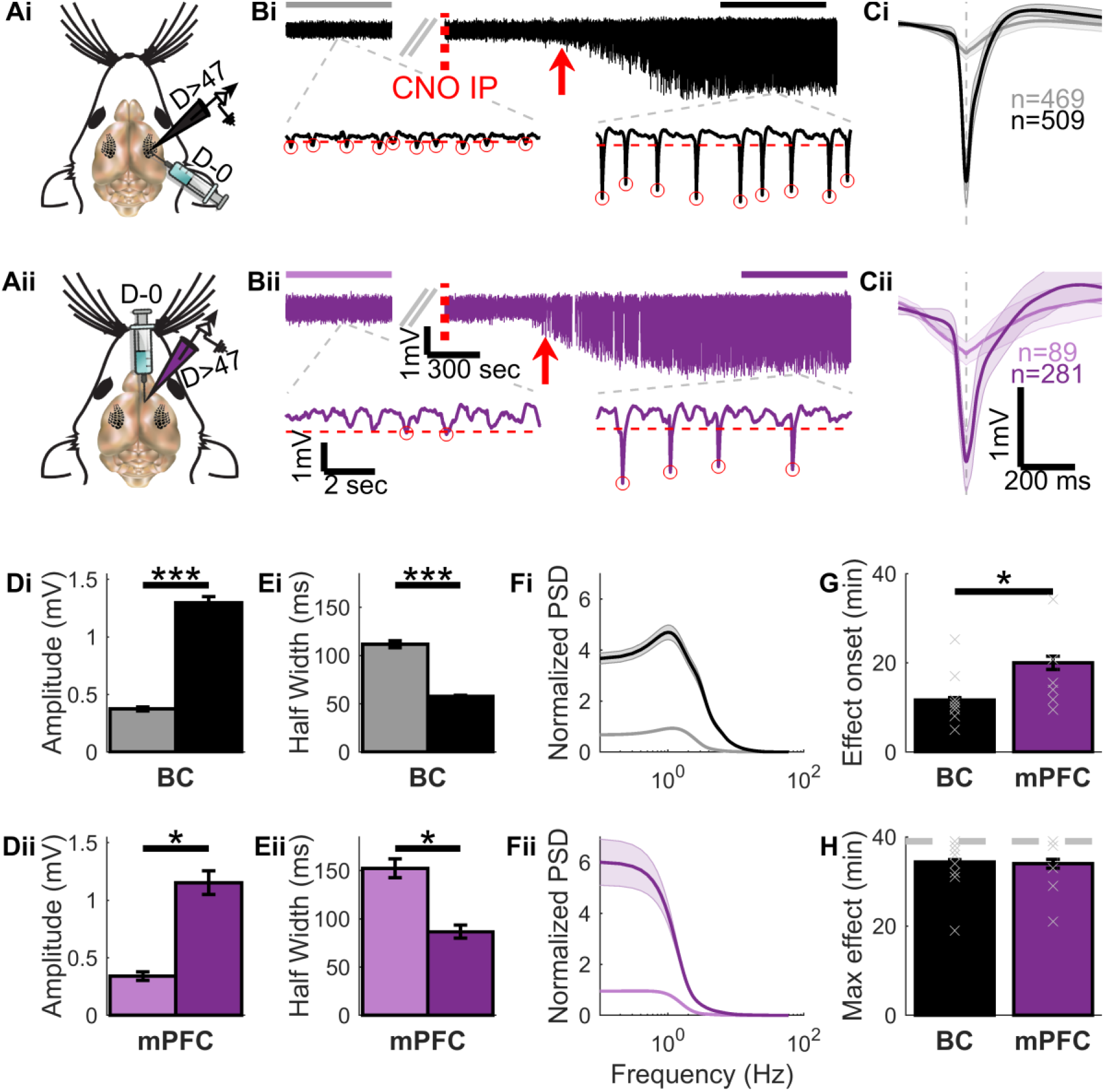
CNO induced epileptiform events in the LFP of anesthetized mice. **A.** Illustration of the experimental design. LFP was recorded at either the BC (Ai) or mPFC (Aii) of anesthetized Gad2-IRES-Cre mice 47 days or more after Cre-dependent hM4D virus was injected to the same site. **B.** Top panels show an example of LFP recorded before and after CNO administration in a mouse injected at the BC (Bi) or the mPFC (Bii). The dashed red line indicates CNO i.p. administration while the red arrow indicates the onset of CNO effect on the LFP. The horizontal lines above the signal mark the ten-minute period used for analysis of baseline (grey/light purple) and post CNO (black/dark purple) signals. Bottom panels show a zoom in of the signal alongside indication of detected events (red circles) and the threshold used in order to detect them (horizontal dashed line). Panels Bi and Bii share the size bars of the top and the bottom panels. Several minutes after the CNO injection large negative peaks started to appear in the LFP of the hM4D expression site regardless of its identity. **C.** Average ± STD of the events detected in the signals shown in panel B during baseline and post CNO at the BC (Ci, grey and black accordingly) and at the mPFC (Cii, light and dark purple accordingly). Panels Ci and Cii share a size bar. The average post CNO event was 5.7 times bigger than the average baseline event in the BC injected mouse, and 3.6 times bigger in the mPFC injected mouse. **D.** Average ± SEM event amplitude across mice injected at the BC (Di) or the mPFC (Dii). The average amplitude of post CNO events was significantly bigger than it was during baseline across mice injected at the BC (N=11, p=0.001) or the mPFC (N=7, p=0.016). **E.** Average ± SEM event half width across mice injected at the BC (Ei) or the mPFC (Eii). The average half width of post CNO events was significantly shorter than it was during baseline across mice injected in the BC (N=11, p=0.001) or the mPFC (N=7, p=0.031). **F.** Average ± SEM normalized power spectrum density (PSD, see methods) at the BC (Fi) and the mPFC (Fii) across mice before and after CNO (grey and light purple vs. black and dark purple accordingly). The maximal PSD following CNO was significantly bigger than baseline PSD at the BC (N=11, p<0.001) as well as at the mPFC (N=7, p=0.008). **G.** Average ± SEM CNO effect onset delay at the BC and mPFC injected mice. On average, CNO started to affect the LFP after 11.7 minutes at the BC and after 20 minutes at the mPFC (p=0.037). **H.** Average ± SEM time to reach maximal CNO effect in the BC and the mPFC (see methods). Around this time the ten-minute window for analysis of post CNO signals was set. This time was analytically constrained to be no longer than 40 minutes after CNO administration (indicated by the dashed grey line). The maximal effect in the signal of many of the mice was close to this limit, regardless of the virus expression site.

**Figure 3.**
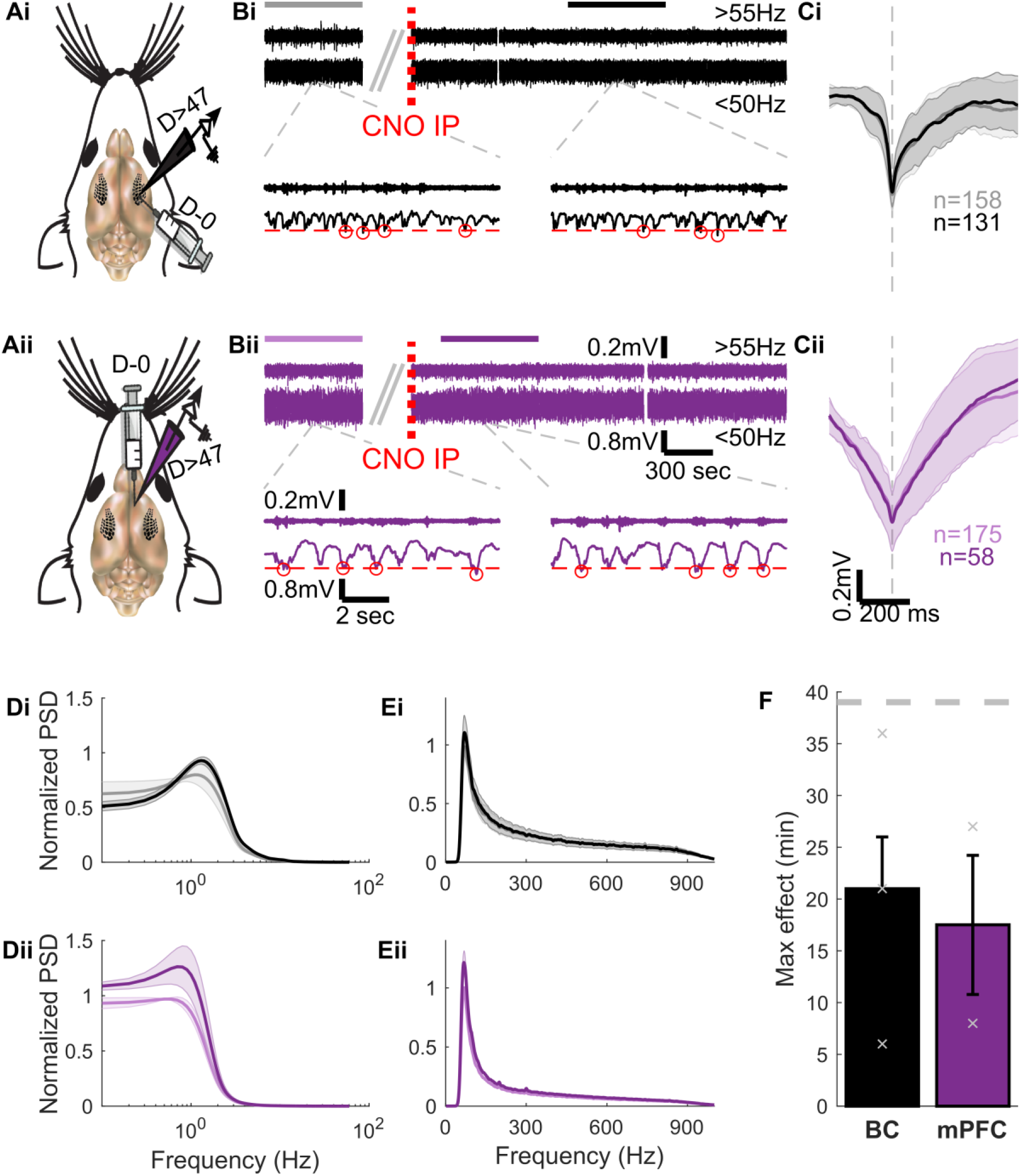
CNO did not induce epileptiform events in the LFP at the absence of hM4D expression. **A.** Illustration of the experimental design. LFP was recorded at either the BC (Ai) or mPFC (Aii) of anesthetized Gad2-IRES-Cre mice 47 days or more after Cre-dependnt virus without the hM4D gene was injected to the same site. **B.** Top panels show an example of LFP and of the high bandpass filtered signal (55-1000Hz, magnified by x4) recorded before and after CNO in a mouse injected at the BC (Bi) or the mPFC (Bii). The dashed red line indicates CNO i.p. administration. The horizontal lines above the signal mark the ten-minute period used for analysis of baseline (grey/light purple) and post CNO (black/dark purple) signals. Bottom panels show a zoom-in of the signal alongside indication of detected events (red circles) and the threshold used in order to detect them (horizontal dashed line). Panels Bi and Bii share the size bars of the top and the bottom panels. No change appeared in the LFP of any of the mice for at least 45 minutes following the CNO injection. **C.** Average ± STD of the events detected in the signals shown in panel B during baseline and post CNO at the BC (Ci, grey and black accordingly) and at the mPFC (Cii, light and dark purple accordingly). Panels Ci and Cii share a size bar. No major difference was apparent between the average events. **D.** Average ± SEM normalized PSD of the LFP signal (<50Hz) at the BC (Di) and the mPFC (Dii) across mice before and after CNO (grey and light purple vs. black and dark purple accordingly). When compared to hM4D expressing mice (see Fig 2F), there was no apparent difference between the PSD of LFP before and after CNO neither at the BC (N=3) nor at the mPFC (N=2). **E.** Average ± SEM PSD of the high pass filtered signal (>55Hz) at the BC (Ei) and the mPFC (Eii) across mice before and after CNO (grey and light purple vs. black and dark purple accordingly). There was no significant difference between the PSD before and after CNO at either site. **F.** Average ± SEM time between CNO injection and reach of maximal LFP variability in a ten-minute window, then chosen for analysis. Within the 45-minute period examined after CNO, all the mice showed maximal variability well before the end of the examined period.

### Propagation of seizure-like activity in-vivo

One key advantage of a focal model with inducible cortical seizures is the ability to examine how local seizure activity propagates to remote areas. We assessed the propagation of CNO-induced epileptiform events by simultaneous recordings at a remote site as well as at the virus injection site. Specifically, we used our system to ask how seizures propagate from primary sensory areas to higher cortical areas, following the bottom-up streams, and vice versa, when they are initiated in higher cortical areas and spread into lower sensory areas.

Towards this aim, we examined how seizures propagate from the primary somatosensory cortex to the mPFC and vice versa. As illustrated in Figure 4A, recordings were acquired from mice injected at the right mPFC 47+ days later at both the right mPFC and the rBC (N=7). Mice injected at the rBC were divided into two groups, one recorded at the rBC and the left BC (lBC, N=5) and the other at the rBC and the ipsilateral mPFC (N=6).

**Figure 4.**
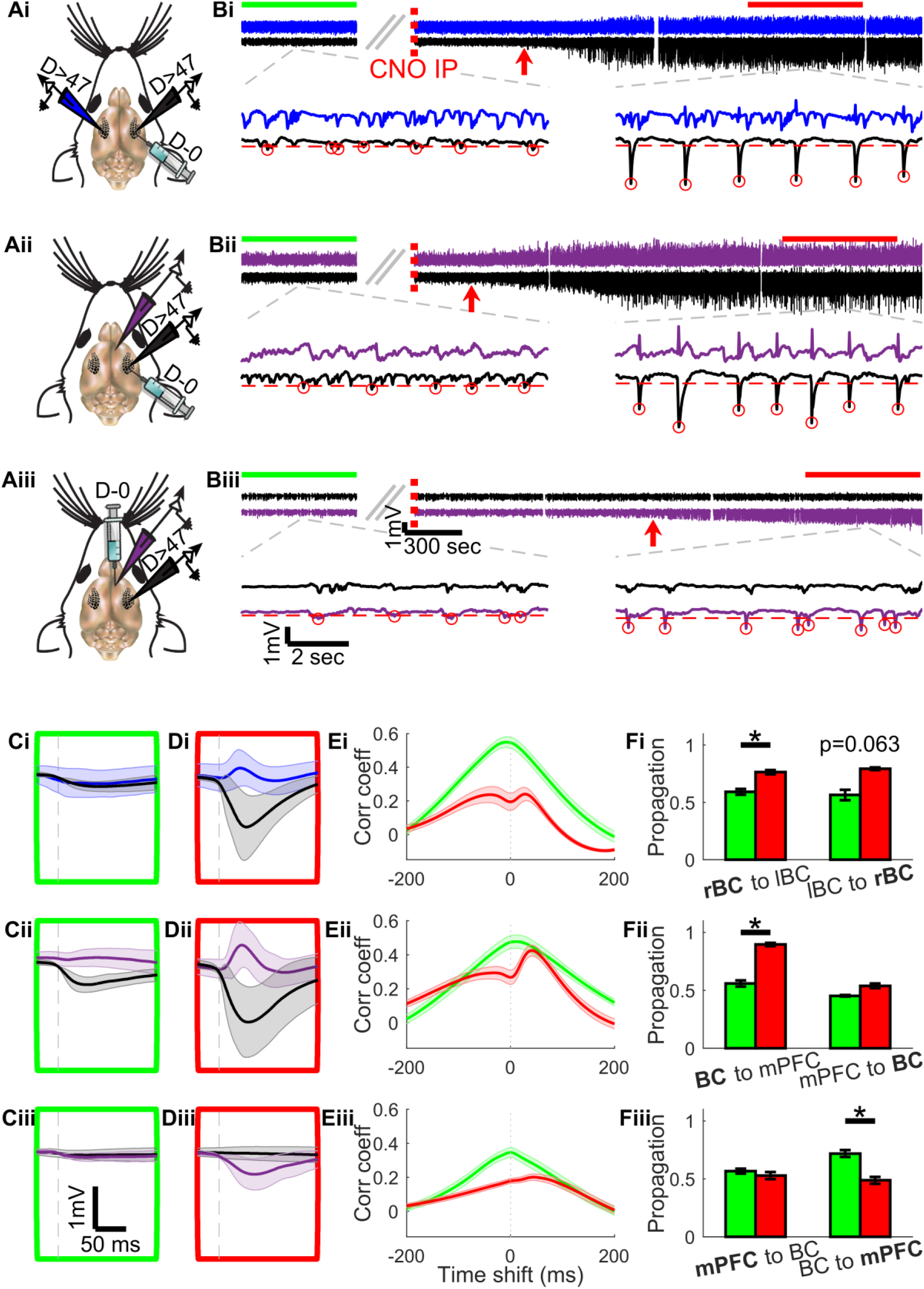
Epileptiform events propagated between certain brain areas. **A.** Illustration of the experimental design. Gad2-IRES-Cre mice were injected at either the BC or mPFC with a Cre-dependent hM4D virus, and at least 47 days later LFP was recorded at the virus injection site as well as one additional area (contralateral BC-i, un-injected mPFC-ii, or un-injected BC-iii). **B.** Top panels show an example of LFP recorded before and after CNO administration in a mouse injected at the BC and recorded in the BC bilaterally (Bi), and in mice recorded at both the mPFC and BC after virus injection to the BC (Bii) or the mPFC (Biii). The dashed red line indicates CNO i.p. administration while the red arrow indicates the onset of CNO effect on the LFP. The horizontal lines above the signal mark the ten-minute period used for analysis of baseline (green) and post CNO (red) signals. Bottom panels show a zoom in of the signal alongside indication of detected events (red circles) and the threshold used in order to detect them (horizontal dashed line). Panels Bi, Bii and Biii share the size bars of the top and the bottom panels. The large negative peaks that appeared following CNO in the injected BC were often accompanied by smaller positive peaks at either the contralateral BC or the mPFC. No such response was observed at recordings from the BC when the virus was injected into the mPFC. **C., D.** Average ± SEM of the average event detected during baseline (C) and post CNO (D) across mice (N=5, N=6 and N=7 for panels i-iii accordingly) aligned by the onset of the virus injection site event. All of panels C,D share size bars. When hM4D was expressed in the BC, about 10ms after the average BC event, a positive peak event was initiated at the contralateral BC and at the mPFC. However, this phenomenon was not observed before CNO, or after CNO when the hM4D was expressed at the mPFC. **E.** Average ± SEM of the cross correlation between the LFP at the hM4D expression site and at the remote recording across mice (N=5, N=6 and N=7 for panels i-iii accordingly) during baseline (green) and post CNO (red). After CNO there is a decrease in the cross correlation around zero for all the mice injected at the BC, but not for the mice injected at the mPFC. **F.** The proportion of events showing significant Granger causality of the virus injection site over the remote site (left) and vice versa (right) during baseline (green) and post CNO (red). There was a significant increase in the percentage of events where the BC as the hM4D expression site significantly Granger caused the remote signal at both the contralateral BC (N=5, p=0.031, one-sided Wilcoxon signed rank test) and at the mPFC (N=6, p=0.016, one-sided Wilcoxon signed rank test). Additionally, there was a significant reduction in the proportion of events where the BC significantly Granger caused the events at the hM4D expressing mPFC following CNO (N=7, p=0.016, two-sided Wilcoxon signed rank test).

Alongside the appearance of epileptiform events at the hM4D expression site, we detected a noticeable change in the LFP of the remote regions in some of the tested expression-remote sites combinations. When hM4D was expressed in the rBC, positive peaks synchronized with the epileptiform events were observed at both the lBC and the mPFC following CNO administration (Fig. 4Bi-ii, the upper trace depicting the recorded activity in the remote site and the trace below it depicts the activity at the injected site). On average, and unlike baseline detected events, a few milliseconds after the onset of almost each rBC post CNO epileptiform event, a positive peak event began at the mPFC and at the lBC, quickly reaching its peak at about 10ms before the average rBC peak (Fig. 4C-D). It is worth noting that volume conductance has been shown to exhibit no jitter between the timing of the activity at the origin and the target sites (Parabucki and Lampl, 2017). However, here the positive peaks at the lBC and at the mPFC were not time-locked with the rBC peaks and showed no correlation with their amplitudes (Fig. 5), thus strongly suggesting that they were not caused by a volume conductance spread.

**Figure 5.**
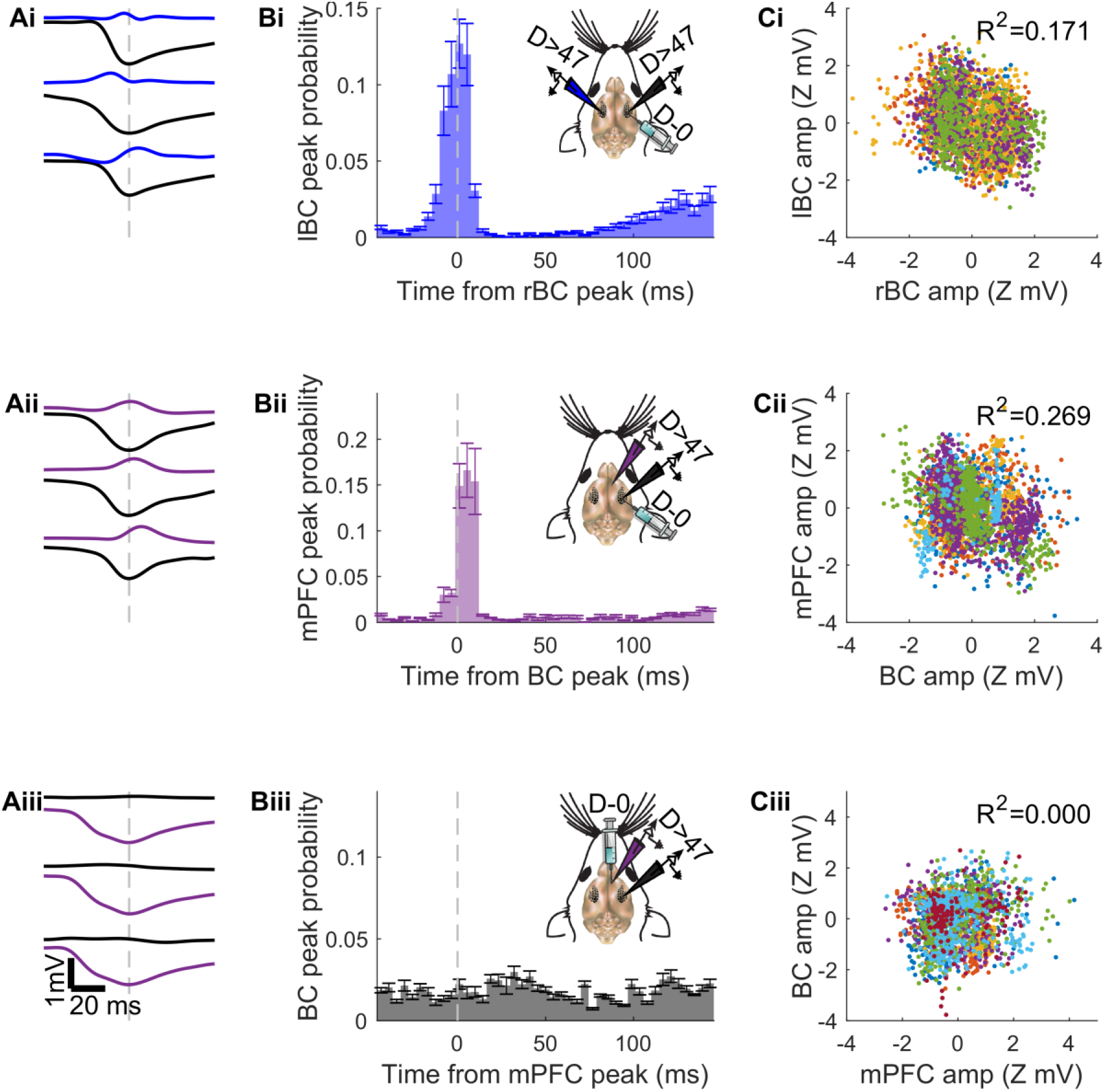
Events recorded remotely from the focus were not caused by volume conduction. **A.** Examples of detected LFP events at the hM4D expression site alongside the LFP recorded simultaneously at the remote site from mice with the following hM4D-remote sites: rBC (black) - lBC (blue, i), rBC – right mPFC (purple, ii), and right mPFC – rBC (iii). Panels Ai, Aii and Aiii share size bars. **B.** Average ± SEM histogram of the probability of certain delays between the remote and the hM4D expression sites event peaks across mice after CNO (N=5, N=6 and N=7 for panels i-iii accordingly). Inserts illustrate the experimental design. In the case of bilateral BC recording, the remote peak alternated between preceding and following the hM4D expression site peak, with a considerable time shift variability within each mouse. In the case of the mPFC as the remote site the peaks often followed the peaks at the hM4D expressing BC but still showed considerable time shift variability per mouse. There was no clear time shift preference for either of the mice injected at the mPFC. **C.** Correlation between post CNO z-scored events amplitudes at the hM4D expression site and at the remote site. Different colored dots represent events recorded at different animals (N=5, N=6, and N=7 for panels i-iii accordingly). For all the three groups tested, the correlation between the amplitude of the events at the hM4D expression and at the remote site was very low (p<0.001, p<0.001, and p=0.346 for panels i-iii accordingly).

The positive peaks at the remote mPFC and lBC also led to the reduction of the overall cross correlation between the LFP signals around lag zero (Fig. 4E). The cross-correlations after CNO injection exhibit a tendency towards double peaks, which can again be explained by the prominent change in the polarity of the events in the remote site.

We also used Granger causality analysis, which provides a statistical measurement of the flow of information between two time series, to find the proportion of events that can be explained by a flow of information from one recording site to the other. There was a significant increase in the proportion of events at the rBC significantly Granger causing the events at both lBC and mPFC following CNO (Fig. 4Fi-ii). Interestingly, there was also a substantial increase in the proportion of events where the lBC signal significantly Granger caused the potentially epileptic rBC signal following CNO, but this increase was not statistically significant. The bidirectionality of the increase in the proportions of significant Granger causing events at the rBC and the lBC is in agreement with the jitter measurement which was centered near zero (Fig. 5Bi). Together these findings suggest that epileptic events can be triggered either by the injected right hemisphere, or by initiation of events in the non-injected side.

Despite the above mentioned propagation of BC events, we found no evidence of epileptiform events propagation from the mPFC to the ipsilateral BC (Fig. 4C-Eiii). On the contrary, a relatively high percentage of the baseline events showed BC Granger causing the signal at the injected mPFC, but this percentage significantly diminished after the CNO was injected (Fig. 4Fiii). All together, these results indicate that seizures initiated in the BC propagate to the mPFC more than seizures that are initiated in the mPFC and examined in the BC.

### High-frequency components of epileptiform activity in-vivo

High frequency oscillations (HFOs) are a prominent hallmark of both human and experimentally induced cortical epilepsies (Jirsch et al., 2006; Zijlmans et al., 2012; Lévesque and Avoli, 2019). We therefore analyzed the high frequency component of the recorded signals (bandpass filtered between 55 and 1000Hz). Encouragingly, we found that the epileptiform events at the hM4D expression site were also accompanied by HFO.

As depicted in the examples shown in Figure 6, across time, the high bandpass filtered signal at the hM4D expression site showed an increase in power after CNO administration, which coincided with the appearance of LFP epileptiform events at the BC (Fig. 6A-Bi) and mPFC (Fig. 6Ci). The increase in power was caused by the emergence of high amplitude HFO events, unobserved before CNO application (Fig. 6A-Cii vs. Fig. 6A-Ciii). These examples also suggest that HFO events only appear in the recordings made in hM4D expressing areas.

**Figure 6.**
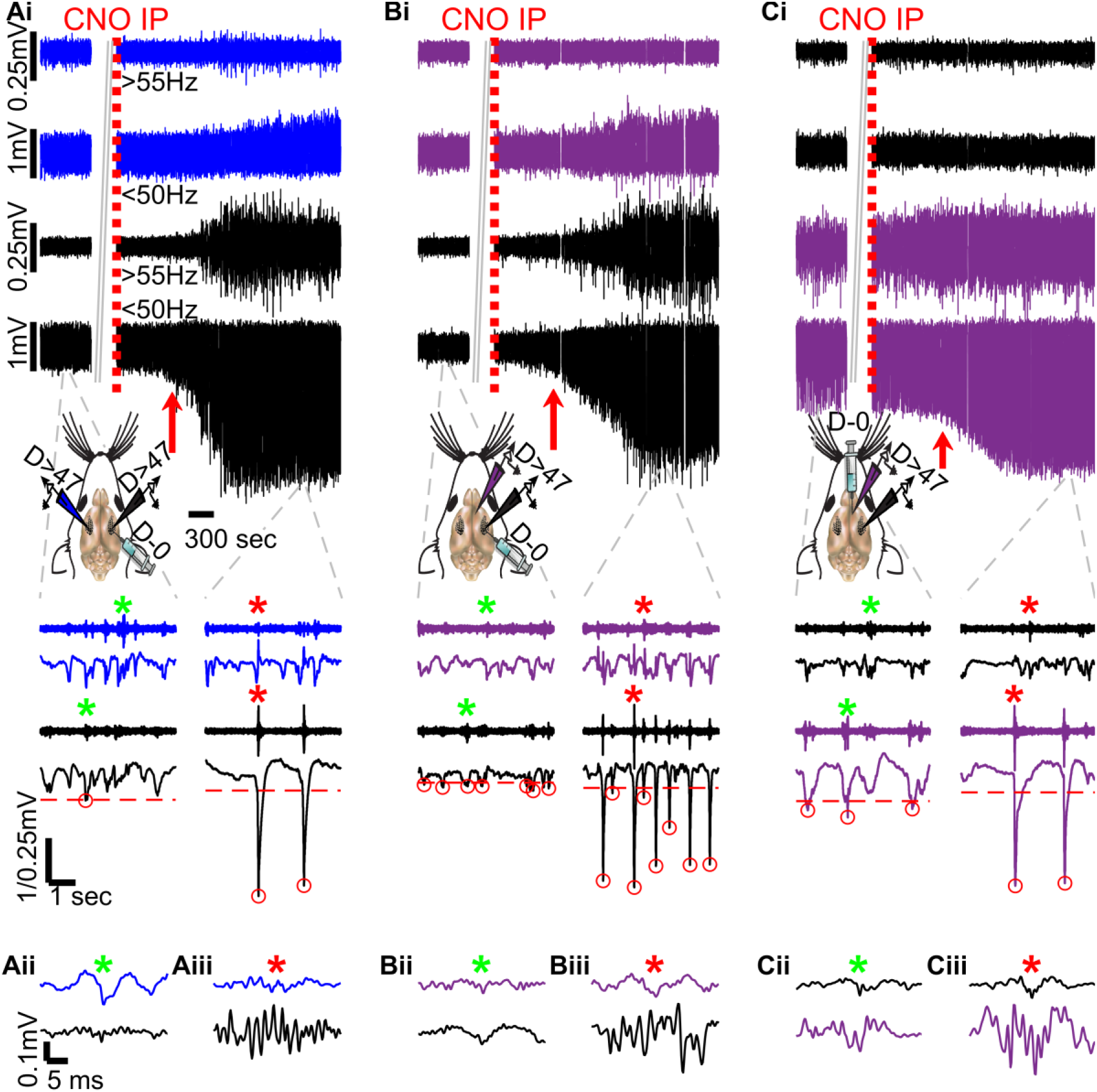
Focus epileptiform events were accompanied by high frequency oscillations (HFO) **Ai.** The top panel shows an example of the LFP and the high bandpass filtered signals (>55Hz, magnified by x4) recorded at the hM4D expressing rBC (black) and the lBC (blue). Inserts illustrate the experimental design. The dashed red line indicates CNO i.p. administration while the red arrow indicates the onset of CNO effect on the LFP. Bottom panels show a zoom in of the signals. **Aii.** A further zoom in on the baseline events in the high bandpass filtered signals marked with green asterisks in Ai. **Aiii.** A similar zoom in on the post CNO events in the high bandpass filtered signals marked with red asterisks in Ai. **B.** Similar to the A subpanels for a mouse expressing hM4D in the BC and recorded simultaneously at the mPFC. **C.** Similar to the A and B subpanels for a mouse expressing hM4D in the mPFC and recorded simultaneously at the BC. The top and bottom panels of Bi and Ci share size bars with their counterpart panels in Ai while panels A-Cii and A-Ciii share size bars as well. All three mice showed an increase in the amplitude of the high bandpass filtered signal at the hM4D expression site following the CNO injection. This increase in power seemed to be composed of big amplitude events, which were highly correlated with the LFP epileptiform events, and encompassed a high frequency oscillation component. However, this effect was restricted to the hM4D expression site in all three mice.

Indeed, multi subject analysis revealed a significant increase in the overall PSD in the ripples spectrum (80-250Hz) and in the fast-ripples spectrum (250-500Hz) for both hM4D expressing BC and hM4D expressing mPFC (Fig. 7A-Ei). While the effect size in the BC was more substantial at the higher frequency band, the opposite was true for mPFC injected mice, leading to a significantly bigger baseline-post CNO change at the mPFC than in the BC at the lower frequency band tested. No significant change was observed in the PSD of any recorded area not expressing hM4D following CNO (Fig. 7A-Eii).

**Figure 7.**
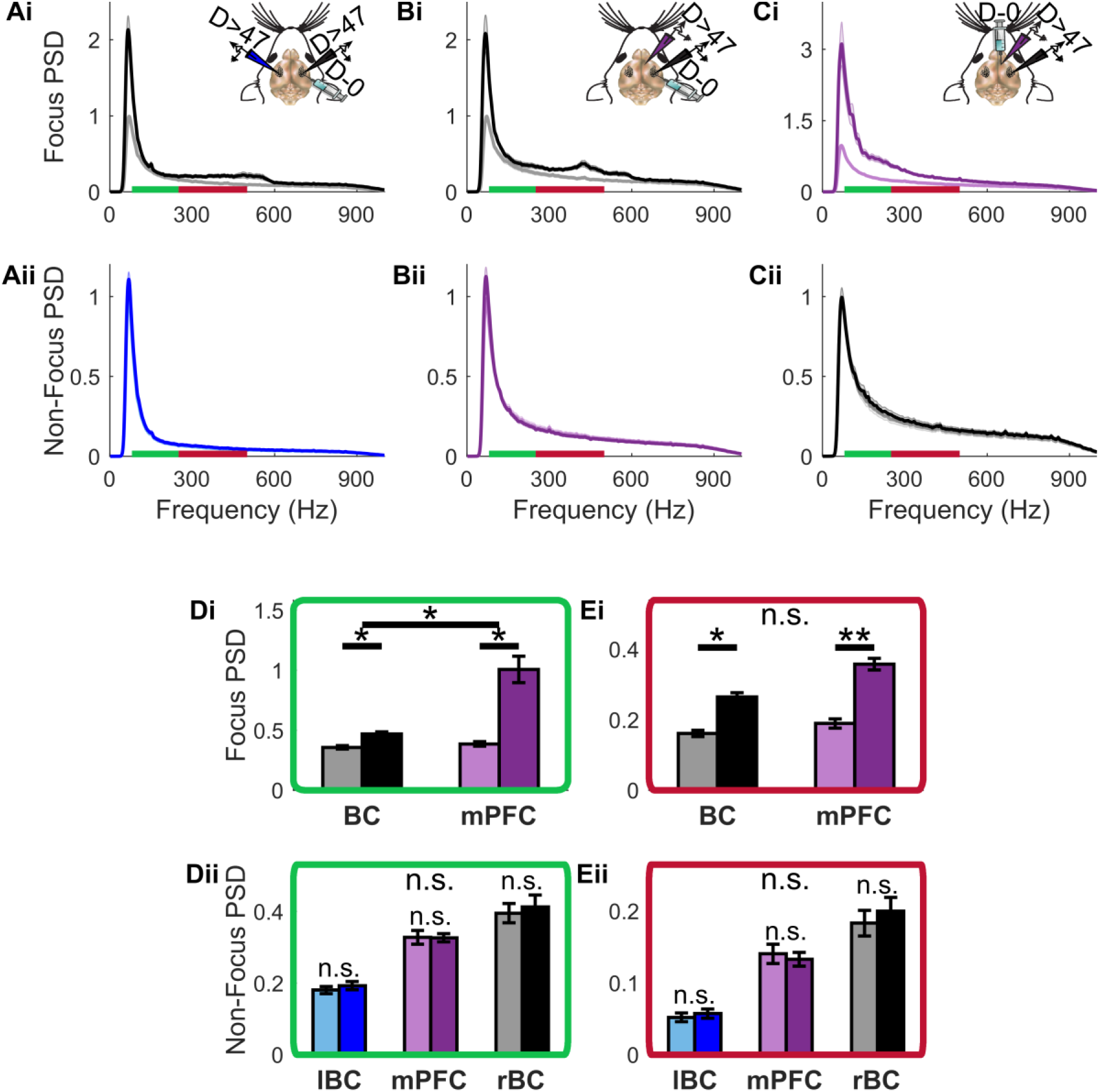
The HFO appearance following CNO was significant across mice. **A.** Average ± SEM normalized PSD of the high bandpass filtered signal (>55Hz) at the hM4D expressing rBC (Ai) and at the simultaneously recorded lBC (Aii) during baseline (grey/light blue correspondingly) and post CNO (black/blue correspondingly) across mice (N=5). The inset illustrates the experimental paradigm, while the green and red lines indicate the analyzed HFO frequency ranges (ripples: 80-250Hz and fast-ripples: 250-500Hz, correspondingly). **B.** Similar to the A subpanels for mice expressing hM4D in the BC and recorded simultaneously at the mPFC (N=6). **C.** Similar to the A and B subpanels for mice expressing hM4D in the mPFC and recorded simultaneously at the BC (N=7). **D.,E.** Average ± SEM of the average ripples PSD (D) and fast-ripples PSD (E) at the hM4D expression site (i, BC N=11, mPFC N=7) and at the remote recording site (ii, BC N=11, mPFC N=7). While the PSD showed no change following CNO at either of the remote sites, there was a significant increase at both expression sites for either one of the examined frequency ranges (ripples: BC p=0.023, mPFC p=0.015; fast-ripples: BC p=0.023, mPFC p=0.002 after Bonferroni correction for repeated measures). Additionally, the baseline to post CNO ripples increase was significantly greater when the hM4D was expressed in the mPFC than in the BC (p= 0.034 after Bonferroni correction).

The increase in overall PSD at the hM4D expression site was most likely due to the contribution of the activity during the epileptiform events, and not of the time between them. This conclusion was supported by similar results arising from the wavelets analysis of the signal around LFP detected events (Fig. 8A-C, Fig. 8Di). Moreover, the wavelet analysis illuminated a post CNO increase in the variability of individual events amplitude across mice regardless of the identity of hM4D expression site and frequency range (Fig. 8Dii). This increase was significant in the BC at both of the tested frequency ranges. In addition, the BC also showed a significant shift in the timing of maximal power at the high frequencies in relations to the LFP peak such that during post CNO events the high frequency activity preceded the LFP peak while the contrary was true during baseline events (Fig. 8Ei). This can be observed in the intensity plots (Fig. 8B-C), where maximal activity appears earlier, during negative time, following CNO application. Lastly, across frequency ranges and regardless of the virus injection site, post CNO events wavelets showed a bigger time lock between their peak and the LFP peak, observed as a reduction in peak timing variability (Fig. 8Eii).

**Figure 8.**
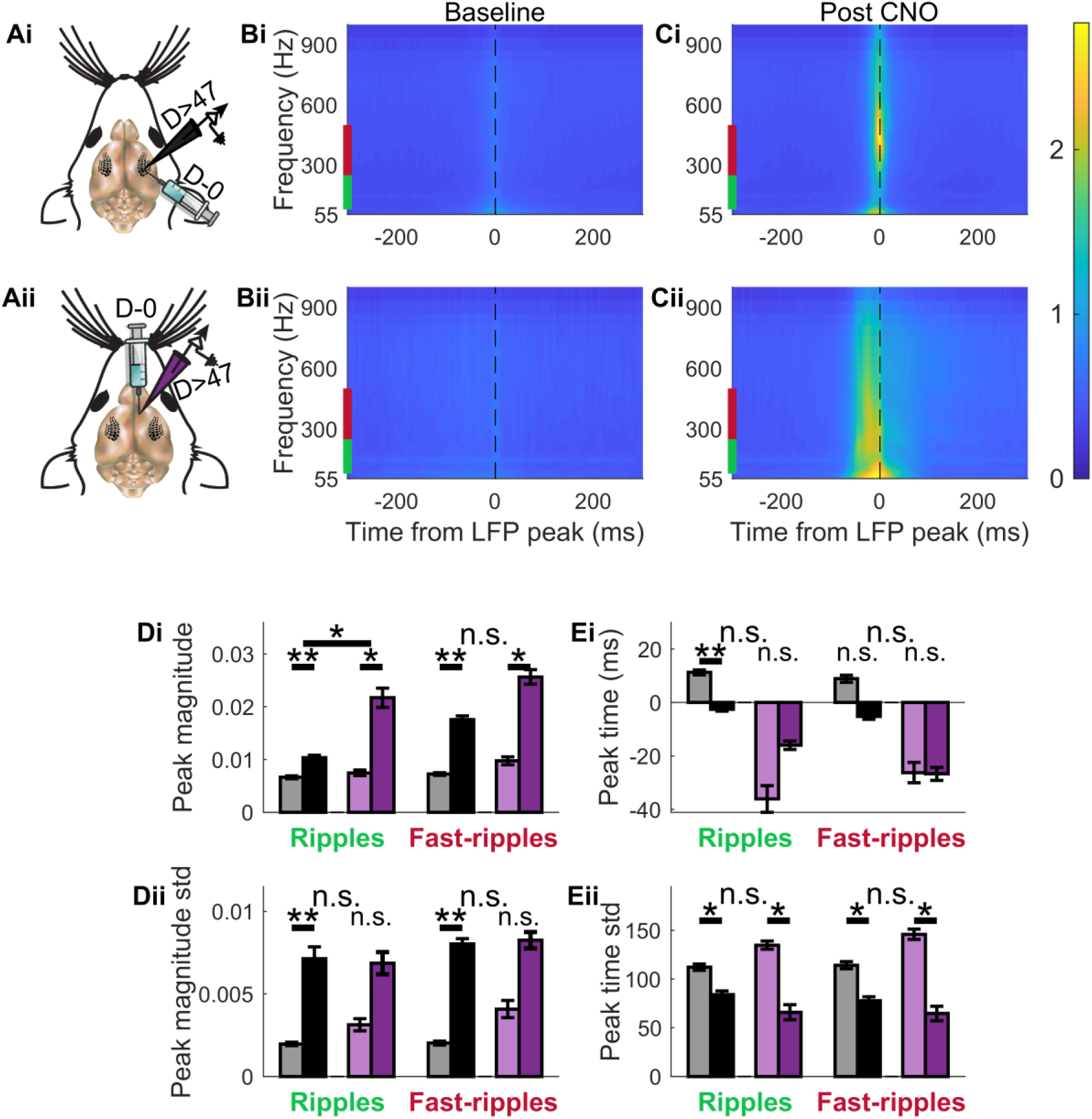
Wavelet analysis of spontaneous focus epileptiform events. **A.** Illustration of the experimental design. **B.,C.** Average of the normalized wavelets of spontaneous baseline (B) and post CNO (C) events across mice injected at the BC (i, N=11) or the mPFC (ii, N=7). **Di.** Average ± SEM maximal wavelet magnitude across mice. For each hM4D expression site and frequency range tested there was a significant increase in the peak magnitude of the events wavelets following CNO (ripples: BC p=0.003, mPFC p=0.047; fast-ripples: BC p=0.003, mPFC p=0.047 after Bonferroni correction for repeated measures). This increase was significantly greater for ripples when the hM4D was expressed in the mPFC than in the BC (p= 0.046 after Bonferroni correction). **Dii.** Average ± SEM wavelet magnitude variability across mice. Following CNO there was an increase in the variability of wavelets magnitudes in all areas and frequency ranges tested. This increase was significant for the BC in both frequency ranges (ripples p=0.003, fast-ripples p=0.003, after Bonferroni correction). **Ei.** Average ± SEM peak wavelet time in comparison to LFP event peak across mice. While baseline wavelets peaks tended to arrive after LFP peaks, following CNO they preceded the LFP peak. This change was significant for ripples (p=0.003 after Bonferroni correction). Wavelet peaks in the mPFC on the other hand preceded the LFP peaks both before and after CNO administration and showed no significant change in the average peak timing. **Eii.** Average ± SEM peak wavelet time variability across mice. The results showed a significant reduction, of similar magnitude, in the wavelets peaks timing variability regardless of the virus expression site identity and frequency range tested (ripples: BC p=0.041, mPFC p=0.047; fast-ripples: BC p=0.020, mPFC p=0.047 after Bonferroni correction for repeated measures).

### Sensory-evoked epileptiform activity

Animal and human studies show that sensory stimulation can trigger epileptic events (Mameniškienė and Wolf, 2018). Examination of the effect of inhibition silencing on the response to sensory stimuli was carried out by providing contralateral whisker pad stimulation to mice expressing hM4D at the BC before and after CNO administration. Beside its effect on the spontaneous activity, we found that CNO also modulated the response to the sensory stimulation. An identical whisker pad stimulation given to the contralateral whisker pad before and after CNO administration induced a far larger response in the BC during the latter (Fig. 9A-B). The response to the stimulus was different in shape and bigger in size in the mPFC and lBC as well following CNO, becoming more similar to the signal shown in these areas during spontaneous rBC epileptiform events. Furthermore, when the stimulus was presented repeatedly at different frequencies after CNO administration there was a reduction in the size of the response to the consecutive stimuli at 3Hz and an even greater reduction at 6Hz, though responses to 1Hz stimuli were all similar (Fig. 9C-E). Control mice lacking the expression of hM4D showed no difference in their whisked pad stimulation response before and after CNO, and only showed minor reduction in size of the response at 6Hz stimulation (Fig. 10).

**Figure 9.**
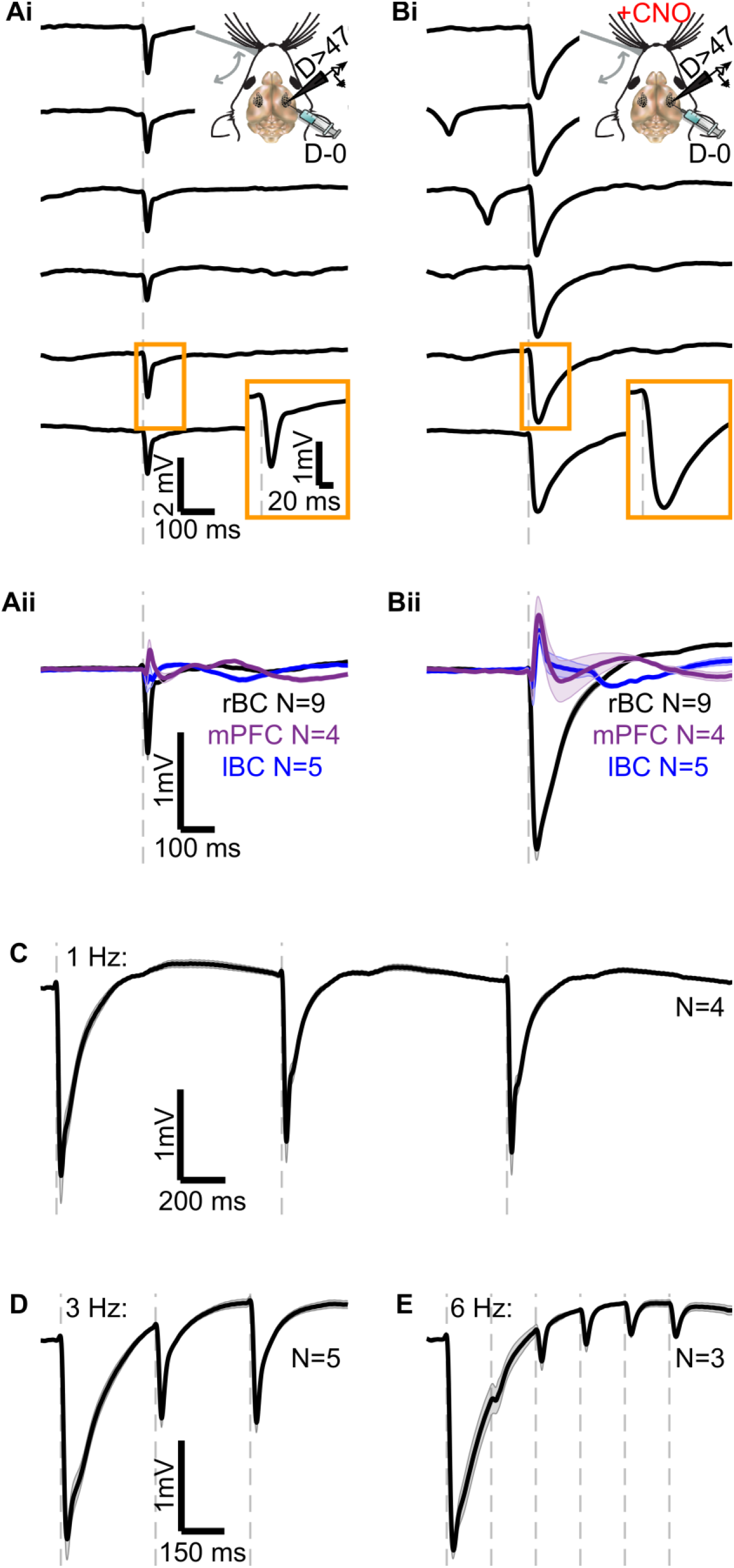
Silencing inhibition using CNO affected the LFP response to sensory stimulation. **Ai.,Bi.** Examples of hM4D expressing BC LFP responses to contralateral whisker pad stimulation before (Ai) and after CNO (Bi). Grey dashed line indicated the stimulation timing. Top insets illustrate the experimental design while the bottom insets show a zoom in of single stimulus responses. Both the overall Ai and Bi panels and their inserts share size bars. The response to the stimulus was rather consistent across trials, but much larger in size following CNO. **Aii.,Bii.** Average ± SEM contralateral whisker pad response across mice at the hM4D expressing BC (black) as well as the mPFC (purple) and the opposite BC (blue) before (Aii) and after CNO (Bii). Panels Aii and Bii share size bars. Not only did the average BC response to the same whisker pad stimulation grew following CNO, but also the mPFC and the contralateral BC average responses grew (and switched polarity in the case of the contralateral BC response). **C., D., E.** Average ± SEM BC response to contralateral whisker pad stimulation at 1Hz (C), 3Hz (D), and 6Hz (E) following CNO. Panels D and E share size bars. While a whisker pad stimulation given at 1Hz always induced an average response similar in size to that of a single post CNO stimulation, repeated stimulations at 3Hz and 6Hz showed a reduction in response magnitude for the latter stimulations.

**Figure 10.**
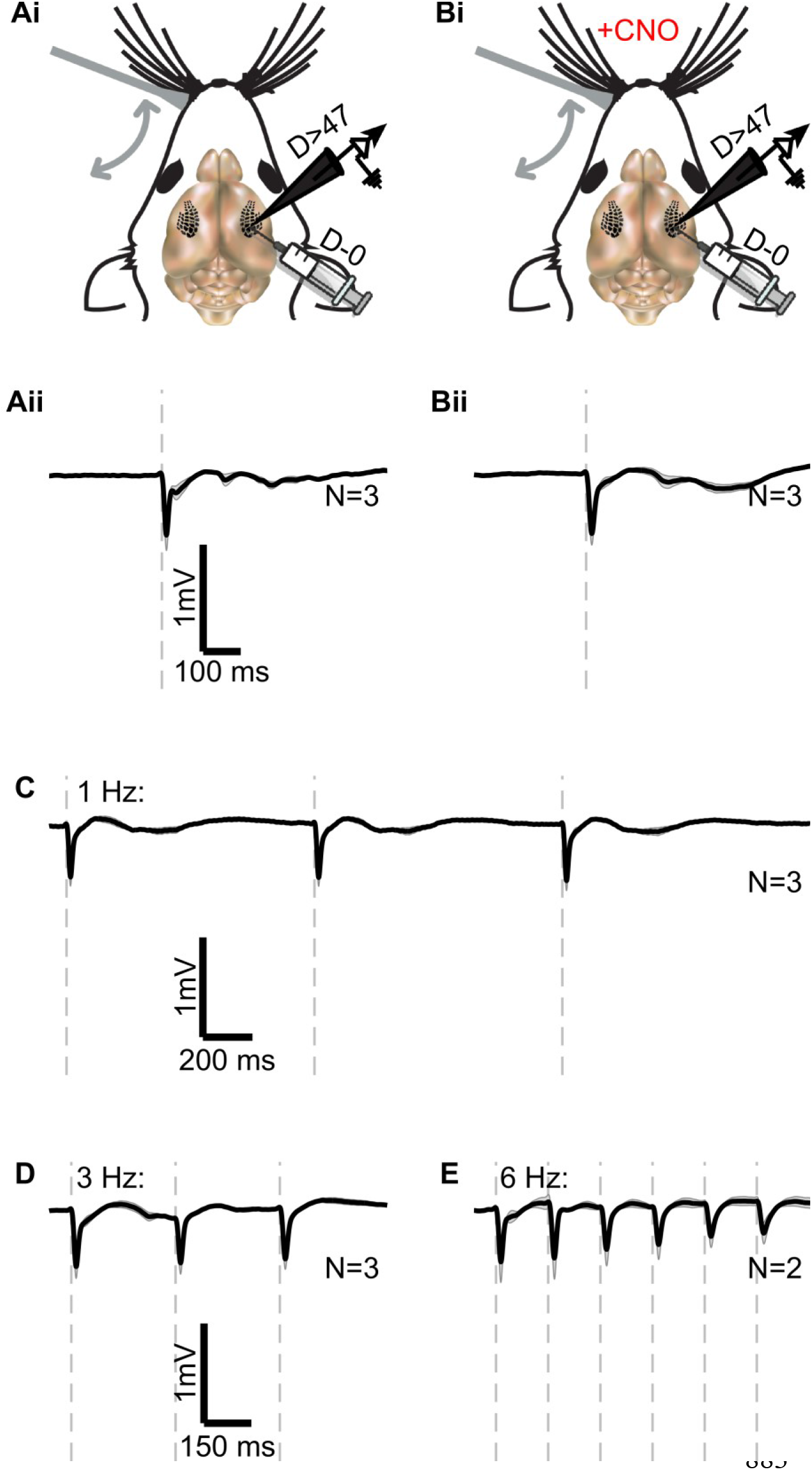
CNO did not affect the LFP response to sensory stimulation in the absence of hM4D. **Ai.,Bi.** Illustration of the experimental design. LFP was recorded at the BC of anesthetized Gad2-IRES-Cre mice 47 days or more after Cre-dependent virus without the hM4D gene was injected to the same site while contralateral whisker pad stimulation took place. The stimulation was presented before (Ai) and at least 45 minutes after CNO administration (Bi). **Aii.,Bii.** Average ± SEM contralateral whisker pad response across mice at the non-hM4D-virus expressing BC before (Aii) and after CNO (Bii). Panels Aii and Bii share size bars. The average response to the stimulation was an effected by the CNO. **C., D., E.** Average ± SEM BC response to contralateral whisker pad stimulation at 1Hz (C), 3Hz (D), and 6Hz (E) following CNO. Panels D and E share size bars. The average response to the stimulation remained similar in shape and size for repeated stimuli given at 1Hz or 3Hz, and only diminished slightly at 6Hz stimulation.

## Discussion

Our findings have shown that targeted chemogenetic inhibition of GABAergic neurons using hM4D can serve as a highly controllable model of focal cortical seizures. Throughout the experiment, CNO reliably induced epileptiform activity within minutes in both awake and anesthetized mice. Reliable focal seizure induction could be achieved in different brain regions, and with unilateral or bilateral expression. We show further that the reliable chemogenetic induction of focal seizures allows a controlled and systematic study of seizure propagation and generalization, exhibiting fundamental characteristics, known in other established animal models of epilepsy and the disease itself.

### Investigation of seizure propagation

Seizure propagation is a major contributor to the clinical symptomatology of epilepsy patients (Nair et al., 2004), and can sometimes predict post-surgical outcomes (Schulz et al., 2000). However, little is known about the mechanisms of seizure spread, why and how they propagate from a certain focus to specific areas and not others. We showed in our model that epileptiform events themselves were restricted to the hM4D expression site. Nonetheless, these events demonstrated indication of propagation from the BC to the mPFC, but not vice versa, and from the BC of one hemisphere to the other (see Fig. 4). Importantly, the propagation could not be explained by volume conductance (see explanation in the results section, Fig. 5).

The propagation speed was assessed by determining the lag between peak activities. In BC-injected mice, the delay measured from BC to mPFC was about 5ms longer compared to the delay from BC to the contralateral BC (Fig. 5). This is consistent with the strong direct BC-BC connection, in contrast to the sparse direct synaptic projections between the BC and the mPFC according to the Allen Mouse Brain Connectivity Atlas (Oh et al., 2014), indicating that seizure propagation likely “hijacks” existing network pathways. We note that anesthesia may be a potential confound in assessing propagation of epileptiform activity, as stronger propagation effects are likely to occur in the absence of anesthesia (Yaffe et al., 2012).

Interestingly, despite direct mPFC-to-BC connections being equally sparse as direct BC-to-mPFC connections, we only observed seizure propagation between these regions when the epileptic focus was at the BC. The selective nature of the propagation signals a stronger bottom-up seizure propagation (i.e. primary sensory area to high cortical area) probability than a top-down one, and it highlights the model’s promise to expand our understanding of this crucial epilepsy property.

### Investigation of sensory driven recruitment into abnormal activity

About 6% of epilepsy patients, with various epilepsy types, report having seizures elicited by external sensory stimuli e.g. sudden noises, flashing lights, tapping and other tactile stimuli (for a review see Trenité, 2012). A review covering a range of sensory-evoked focal seizures emphasized the anatomical epileptic focus and its abnormal neuronal activity as key predictors of sensory triggers (Mameniškienė and Wolf, 2018). In accordance with this statement, our results demonstrated a change in the response to whisker-pad stimulation under the influence of CNO (Fig. 9) which required pre-existing hM4D expression (Fig. 10). The post CNO response was similar in shape and size to that of spontaneous epileptiform events, both at the epileptic BC and at remote regions, suggesting a possible triggering of the propagating epileptiform events by innate network inputs.

Our model thus also allows an examination of focal sensory-induced ictogenesis and network characteristics. This is demonstrated by our observation that the massive network recruitment that is needed to induce an epileptiform event has a refractory-like period preventing the sensory triggering of a full-size event within 0.33-1 seconds from a previous sensory-evoked event (Fig. 9C-D).

### Properties of focal epileptiform activity and HFOs

The CNO evoked epileptiform events in our experiments were also accompanied by HFOs (Figures 6–8). The existence of HFOs is regarded as a reflection of a dysfunctional neural network (Jefferys et al., 2012) and is highly correlated with human and animal seizures alike (Jirsch et al., 2006; Zijlmans et al., 2012; Lévesque and Avoli, 2019). HFOs are an important interictal marker of focal epileptogenic zones, and have been clinically used to localize such areas (Bragin et al., 1999; Andrade-Valenca et al., 2011). Importantly, several reports convergently describe that removal of HFO-generating areas leads to improved surgical outcomes (Akiyama et al., 2011; Haegelen et al., 2013; Weiss et al., 2013; Kerber et al., 2014). Though it has largely been referred to as an interictal phenomena, HFO components are also found during ictal and preictal periods (Jirsch et al., 2006; Jacobs et al., 2009). Taken together, it is evident that our model shares a key characteristic of focal epilepsy onsets in humans.

Our analysis revealed the emergence of both ripples (80-250Hz) and fast ripples (250-500Hz) during the CNO induced seizure at either focus site (Figures 7–8). While the difference between the mechanisms governing the two phenomena are thoroughly discussed elsewhere (Jefferys et al., 2012; Lévesque and Avoli, 2019), it is worth noting that ripples have predominantly been associated with inhibitory postsynaptic potentials (Buzsaki et al., 1992; Ylinen et al., 1995; Chrobak and Buzsáki, 1996; Klausberger et al., 2004; Klausberger and Somogyi, 2008). The presence of ripples during massive silencing of the local GABAergic cells observed here implies there is at least one additional cause to epileptic ripples.

### Discussion of the chemogenetic model in relation to other models of focal cortical epilepsy

While common seizure models have mainly targeted the hippocampus, several studies have demonstrated control over seizure induction at cortical areas. Such studies traditionally utilized either intracranial chemoconvulsant injections (Takebayashi et al., 2007), or repetitive electrical stimulation (Collins et al., 1983). With the development of stronger genetic-based manipulation tools, new ways to control seizure induction came to light in the form of chemogenetics and optogenetics.

To the best of our knowledge, though many chemogenetic endeavors have been devoted to seizure suppression (see Forcelli, 2017, for a fairly recent review), only a few chemogenetic seizure models have been suggested, and of which, none has featured localized seizure foci. Several optogenetic studies, on the other hand, have shown seizure inducibility at restricted cortical areas.

When used alongside known epilepsy models, optogenetic neuronal activation has been shown to trigger seizures in the somatosensory cortex of WAG/Rij rats (Wagner et al., 2014), and in the entorhinal cortex using the *in-vitro* 4-aminopyridine epilepsy model (Shiri et al., 2015, 2016). In addition, optogenetic activation of pyramidal cells in the primary motor cortex of naïve mice was shown to induce epileptiform activity if given repeatedly (Cela et al., 2019) or at incrementally increasing intensities (Khoshkhoo et al., 2017).

Similarly to these models, our model allows researchers to trigger seizures upon demand at a cortical site of their desire. In contrast, our model achieves this goal by selective chemogenetic inhibition of GABAergic cells, rather than using genetic manipulation in order to activate neurons.

We believe that silencing inhibition via CNO administration to Gad2-IRES-Cre mice intracranially injected with Cre-depended hM4D virus can be used as a seizure model that possesses many crucial features for experimental use. This model is highly reproducible and allows excellent control over seizure onset. It encompasses proven versatility in the location and size of the seizure focus site, thus allowing scientists to study either convulsive or non-convulsive seizures, generalized or partial seizures, etc. It is relatively inexpensive, can be used across consciousness states, and other than the initial virus injection, it does not require complicated or invasive procedures given that CNO is orally bioavailable (Bender et al., 1994). Along with the experimental appeal of this method, it holds a strong resemblance to human seizures in propagation, sensory triggering and oscillatory activity, and therefore embodies great promise as a potential testing model for focal epilepsy treatments. Future work may test known anti-convulsant medications using this model to determine whether it may be suitable for testing possible treatments for drug-resistant epilepsy. Additionally, this model may be used alongside modern neuroscience methods, such as cell-type-specific neuronal imaging, to better characterize seizure spread along different cortical pathways.

## Acknowledgments

Ilan Lampl is the incumbent of the Norman and Helen Asher Professorial Chair at the Weizmann Instituteof Science. Yonatan Katz is incumbent of the Marianne Manoville Beck Research Fellow Chair in Brain Research. This research was supported by DFG (SFB 1089) to HB and IL, the DFG (BE 1822/11) to HB, EraNet (DeCipher Neuron 01EW1606) to HB and IL, and Israel Science Foundation (ISF 1539/17) to IL. This research was also supported by Marianne Manoville Beck Laboratory for Research. We thank all the members of the I.L. laboratory and especially Rebekah Tumasus, Ana Parabucki, and Yael Oran for their helpful contributions.

